# Subtle Structural Translation Remarkably Modulates the Super-Resolution Imaging of Self-blinking Rhodamines

**DOI:** 10.1101/2022.11.20.517287

**Authors:** Ying Zheng, Zhiwei Ye, Yi Xiao

**Affiliations:** State Key Laboratory of Fine Chemicals, Frontiers Science Center for Smart Materials Oriented Chemical Engineering, Dalian University of Technology, Dalian 116024, China

## Abstract

The evolution of super-resolution imaging techniques is benefited from the ongoing competition for optimal rhodamine fluorophores. Yet, it seems blinded to select the best one among different rhodamine derivatives for specific labeling and imaging, without the knowledge on imaging impact of even the minimum structural transform. Herein, we have designed a pair of self-blinking sulforhodamines (STMR, SRhB) with the bare distinction of methyl or ethyl substituents, and engineered them with Halo protein ligands. Although the two present similar spectral properties (λ_ab_, λ_fl_, □, etc.), they demonstrated unique single-molecule characteristics preferring to individual imaging applications. Experimentally, STMR with high emissive rates was qualified for imaging structures with rapid dynamics (endoplasmic reticulum, mitochondria), and SRhB with prolonged on-times and photostability was suited for relatively “static” nuclei and microtubules. Utilized this new knowledge, the mitochondrial morphology during apoptosis and ferroptosis was first super-resolved by STMR. Our study highlights the significance of even the smallest structural modification to the modulation of super-resolution imaging performance, and would provide insight for future fluorophore design.

## Introduction

Super-resolution microscopies have transformed the landscape of biological fields by providing unprecedented resolution beyond Abbe diffraction limit.^1–4^ These microscopies demand fluorophores with robust brightness, outstanding photostability and specific labeling functionality, thus transforming the fluorophore engineering into a systematic work.^5–8^ Yet, the engineering work is challenged by the knowledge gap on the impact from molecular structural adjustment to the modulation of super-resolution imaging performance, especially for single-molecule localization super-resolution microscopy (SMLM).^6^ The localizing in this microscopy is a process of sparse sensing and recovering of molecular positions in the spatial and temporal domain, and is sensitive to the perturbations of emissive and non-emissive states from any structural changes.^9–12^ Thus, it is a necessity to investigate any relevant impacts stepping from the minimum group insertion.

Rhodamine is the adequate fluorophore scaffold for super-resolution imaging credited to their robust photophysical properties,^13–21^ and self-blinking possibilities from thermo spirocyclization equilibrium.^22^ In a landmark work, Urano et al. first introduced a self-blinking rhodamine (HMSiR) for SMLM imaging through optimizing of intramolecular nucleophilic fluorophores.^23^ Lavis et al. and Xiao et al. utilized steric hinderance and electronic appealing effect to suppress the twisted intramolecular charge transfer (TICT) effect, and increased the brightness for SMLM imaging.^24^ Hell et al. designed and synthesized a series of N,N’-di-tert-alkylrhodamines to overcome the spectral blue-shift caused by the photobluning.^15^ Johnsson et al. introduced different electron-withdrawing groups (EWGs) to the amide units, to increasing the cell permeability and reducing non-specific background signal, and achieved wash-free super-resolution imaging.^25^ Xu et al. replaced the benzene ring of rhodamine with 1,3-disubstituted imidazole, which increased the scintillation ability of rhodamine in thiol imaging solution, making it beneficial for two-color super-resolution imaging.^26^ Although these reports suggested a diversity of structural modifications to solidly improve the super-resolution imaging capabilities, it could hardly predict the imaging impact from any future substitution changes up to now. Hence, rhodamine is selected in this study to investigate its imaging modulation impact induced by structural transforms.

Tetramethyl rhodamine (TMR) and rhodamine B (RhB) constitute an efficacious pair to investigate the impact of minimum substituents.^27, 28^ The two are structurally homologous to each other, with the only distinction of arylamine groups (ethyl for RhB, methyl for TMR). RhB and TMR exhibits similar bulk spectral properties (λ_ab_\λ_fl_\□ in ethanol with 0.1% TFA: 553\578\0.52, TMR; 548\574\0.55, RhB; the unit of λ is nm).^15, 29–31^ As a result, it is reasonable to infer that the two fluorophores should have similar imaging capabilities. Indeed, they are equally qualified for conventional/confocal imaging, and both of their analogues have been adopted in SMLM imaging by different research teams, confirming their general applicability.^14, 19, 25, 32–34^ It is surprising that the utility frequency in these reports implied a privilege of TMR over RhB in super-resolution imaging. Yet, it is a mist to tell whether this privilege is a prejudice as the existing reports were conducted on different microscopies under various labeling conditions. Thus, it is worthwhile to uncover their single-molecule fluorescent differences from a “pure” perspective of their molecular structure distinctions, to maximizing their utilities in super-resolution imaging.

To rigorously uncover the mist, two prerequisites should be satisfied: identical labeling with accuracy and self-blinking to remove controversial issues of utilizing self-optimized imaging buffers. The self-labeling protein tag would be a rational choice, as it is facilely realized the same protein ligand for different fluorophores and it provides identical protein microenvironments through binding target fluorophores. The self-blinking properties might be introduced through the adjustment of nucleophilic ability of the amide group in the spiro ring.

Herein, we designed and synthesized sulfonamide derivatives of tetramethyl rhodamines (STMR) and rhodamine B (SRhB), to introduce self-blinking mechanics into the rhodamines, and engineered them with Halo-tag self-labeling protein technology, to explore the imaging impact of methyl or ethyl substitutions (Figure 1a). Single-molecule experiments revealed a large quantity of switching events for STMR and SRhB, validating their self-blinking functionality, yet their single-molecule events demonstrated unique brightness, on-time, and photostability characteristics. The disparity suggested their individual imaging suitability, which was experimentally verified in super-resolution imaging of multiple organelles in living cells. With the preference suggestion from the new discoveries, the dynamics of mitochondrial morphology during apoptosis and ferroptosis were successfully reconstructed with STMR.

**Figure 1.**
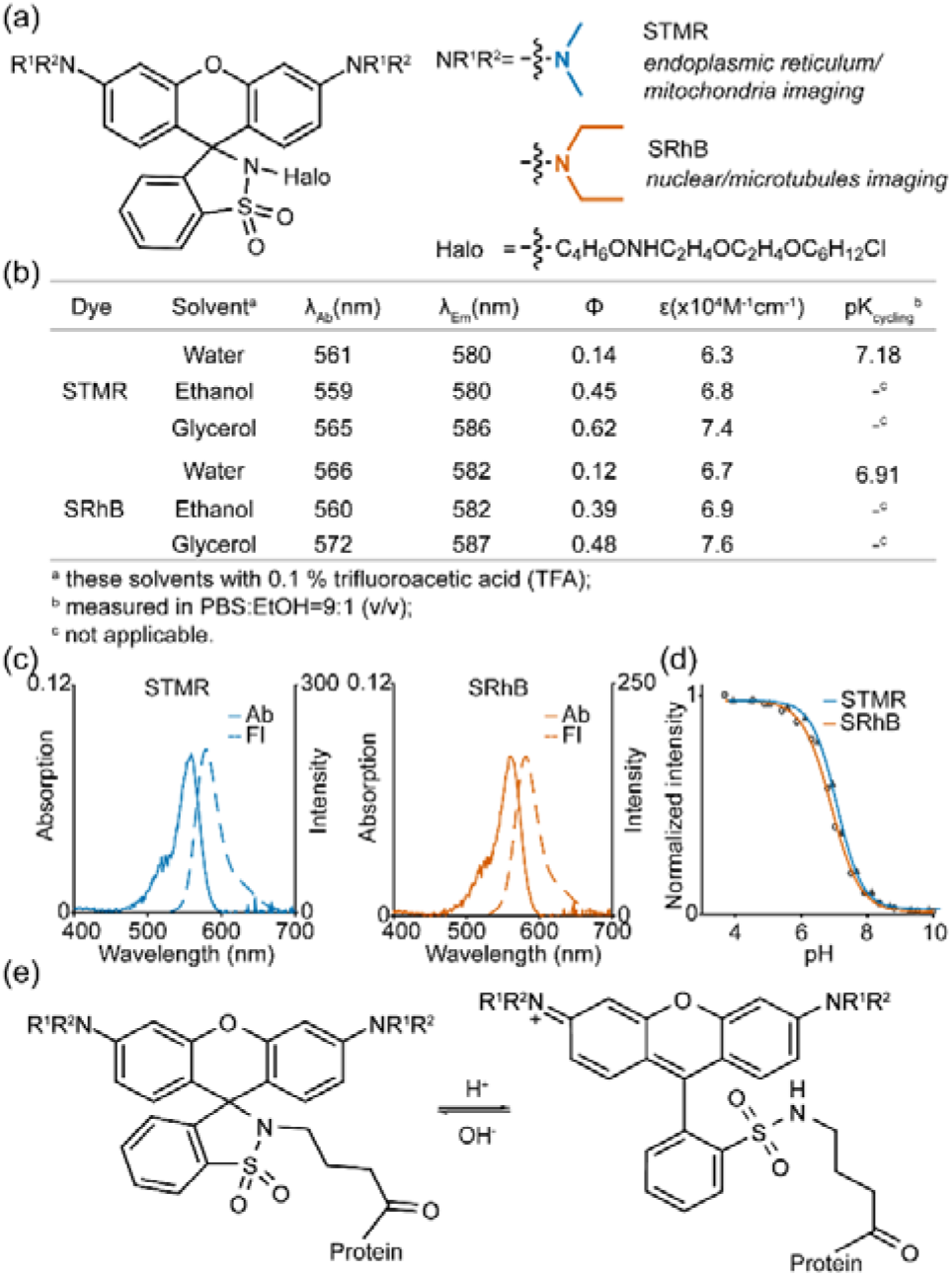
(a) The molecular structure of SRhB and STMR. (b) Summarized spectroscopic properties. (c) Absorption and fluorescence spectra of STMR and SRhB, respectively. (d) Integrated emission intensity vs pH of both STMR and SRhB in C_2_H_5_OH/PBS (v/v = 1:9). (e) Spontaneous spirocyclization equilibrium of sulforhodamine.

## Results and Discussion

### Molecular Design

To analyze the impact from the subtle structural adjustment, the prior need is the switching performance of fluorophores.^9, 35^ Conventional rhodamine dyes mostly require photoactivation or imaging enhancing buffers to achieve fluorescent sparsity.^36–38^ Such complexity might make misjudge in understanding the structural effect. This issue could be addressed with the development of self-blinking rhodamines with thermo-driven spirocyclization equilibrium, which was first introduced by Urano in a landmark work,^23^ and followingly widely utilized for SMLM imaging.^16, 39–41^ Yet, conventional self-blinkable rhodamines constructed on silicon rhodamine emit far red fluorescence, and are not available for other spectral regions. In green region, sulforhodamine has raised our interests as it also possesses spirocyclization equilibrium. Unfortunately, sulforhodamine has been previously considered inapplicable to work as a self-blinking fluorophore in SMLM imaging, as its higher p*K*_cycling_ (~7.0) may result in a too large portion of ring-open molecules at physiological pH to meet the requirement on emission sparsity.^23^ However, we have recently confirmed this concern is not fully justified. We found the fast self-blinking of sulforhodamine is even better than classical silicon rhodamine (HMSiR), and confirmed its applicability in live cell SMLM imaging. The good performances of sulforhodamine is ascribed to its larger recruiting rate, a key kinetics parameter independent on excitation intensity.^42^ Therefore, to broaden our understanding on this new type of self-blinking fluorophores, we prepared two derivatives, STMR and SRhB (Figure 1a) for the comparison study on the imaging impact from methyl and ethyl substituents.

Labeling is the other control factor for our investigation. It is challenging to eliminate the concerns of microenvironment and linking groups, especially when the targeting-unit binding of two fluorophores are independently optimized to suit cross-linked organelles with dynamic environment in living cells. This issue could be addressed by self-labeling protein technology, which enables fluorophores with identical engineering to covalently label to any proteins of interest.^43–46^ Moreover, the same structural protein could be deployed to study the imaging impact from the subtle structural change. Therefore, both STMR and SRhB is engineered with Halo-tag ligands to investigate their super-resolution imaging properties at versatile targets. To further remove the redundancy of cell variability in transient gene expression, single cell with stable expression was screened, cultured and utilized for our study.

### Spectral results

We first investigated the spectral properties of STMR and SRhB. The visible absorption (λ_ab_ = 559-572 nm) and fluorescence (λ_fl_ = 580-587 nm) are similar for the two fluorophores in various solvents (including 0.1% TFA) (Figure 1b). For the peak wavelength, both λ_ab_ and λ_fl_ of SRhB are slightly red-shift 1-7 nm compared to that of STMR, owing to the enhancement of electron donor effect of amine substituents. The fluorescence quantum yields (□) of STMR and SRhB are weak in aqueous solution and significantly enhance in viscous solvent (water/ethanol/glycerol: 0.14/0.45/0.62, STMR *vs* 0.12/0.39/0.48, SRhB), in which the torsion of amine substituents might be suppressed. Moreover, STMR and SRhB exhibited proximate spirocyclization process, with similar p*K*_cycling_ values and cycling equilibrium rates (7.1, STMR *vs* 6.9, SRhB Figure 1d and 1e). The spectra of the two fluorophores demonstrate high similarity, due to their identical scaffolds.

### Single-Molecule Photophysics

The single-molecule photophysics of STMR and SRhB were investigated in vitro (data of STMR has also been utilized in our anther study^42^ to explain the self-blinking). Fluorophores were bounded to Halo proteins, allowing studies to perform in phosphate buffered saline (PBS) to mimic the physiological labeling condition. A set of laser powers were deployed to depict the full spectrum of single-molecule characteristics of the two fluorophores (data at 2 kW/cm^2^ were reported, unless stated otherwise). The fluorophores were sparsely dispersed in the field to avoid signal overlapping, as evidenced in the max-projection views in Figure 2a and 2b. Typical molecular fluorescent trajectories reveal fast transition behaviors for the two fluorophores, suggesting the existence of self-blinking characteristics. Although the bulk fluorescence of STMR and SRhB were similar, their single-molecule fluorescent events shown uniqueness. Statistically, STMR exhibited high single-molecule brightness (371 vs 319; order in STMR, SRhB, following the same; Figure 2c) whereas SRhB presented prolonged on time (25 ms *vs* 64 ms; Figure 2d), and photostability (4.2 s *vs* 8.1 s; Figure 2e) regardless of the excitation powers. The brightness of STMR might be reasoned to its slightly higher quantum yields compared to SRhB and the photostability results suggest ethyl substitutions enhance the stability of rhodamine fluorophore. Owing to its high brightness, STMR demonstrates a slightly higher localization precision (13.12 *vs* 13.51; 10 ms acquisition time) than SRhB (Figure 2f). Figure 2g summarizes the single-molecule characteristics. Despite of their different substituents, the two fluorophores have fast recruiting rates (also remarked as turn-on rate, see study[x]; k_rc_ = 0.372 s^−1^ *vs* 0.313 s^−1^) and large switching numbers (19.0 *vs* 14.0), met the standard as a self-blinking fluorophore for SMLM imaging. In summary, the slight transfer from methyl to ethyl does not interfere the self-blinking characteristics of STMR and SRhB, but provides unique single-molecule fluorescent events with different brightness, on time and photostability.

**Figure 2.**
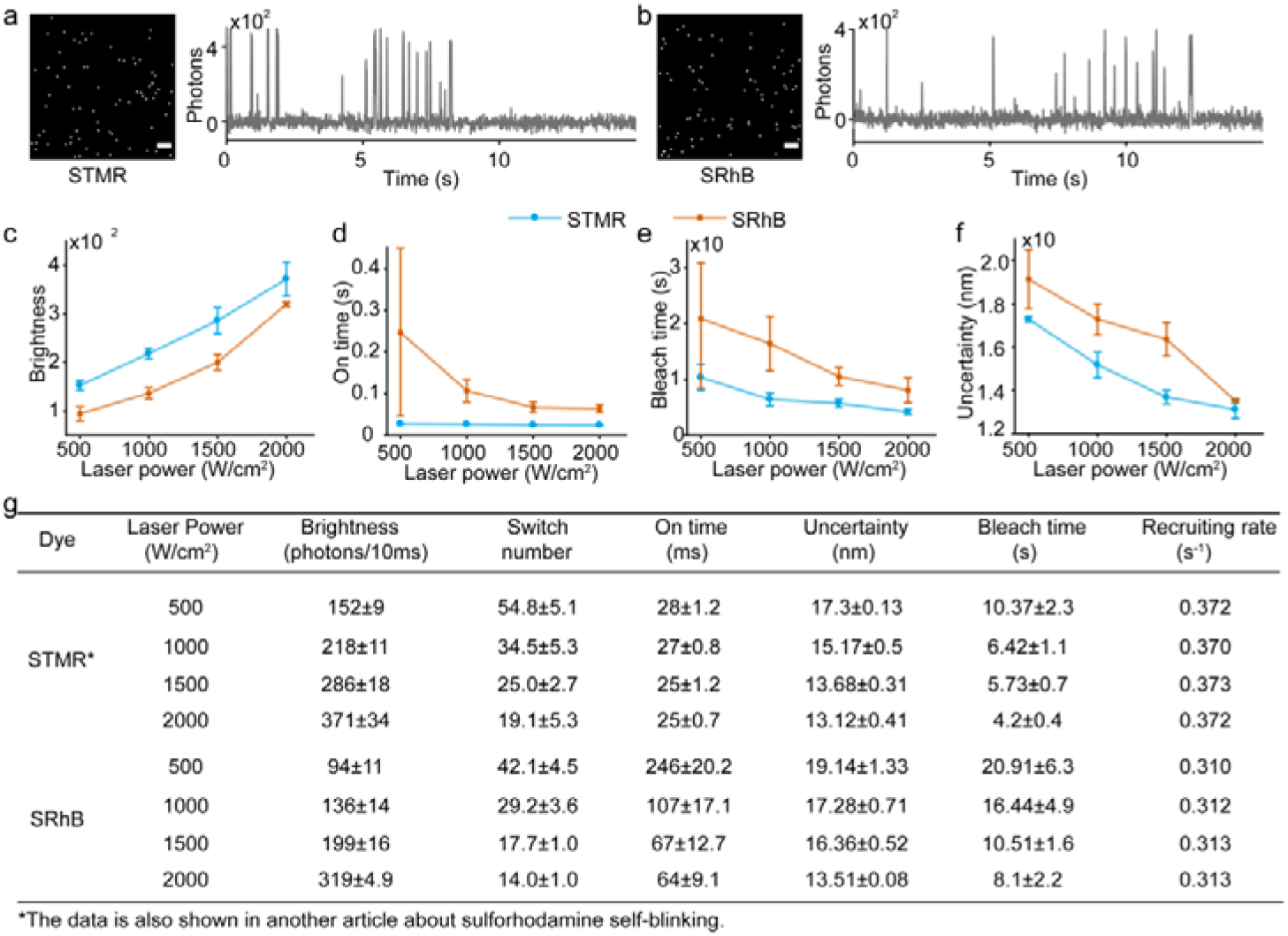
Single-molecule photophysical of STMR and SRhB bounded to Halo proteins in PBS under a set of irradiation powers. (a, b) Max projection view of imaging stacks shows the sparse distributed single-molecule fluorescent signals and typical fluorescence trajectories of two rhodamines under continuous irradiation of a 2 kW/cm^2^ laser. The single-molecule brightness (c), the on-time (d), the bleach time (e), and uncertainty (f) as a function of laser power. The summary of single-molecule characteristics (g) of two rhodamines under continuous irradiation with different laser power. Scale bars: 4 μm.

For living-cell imaging applications, it could be inferred that STMR is suitable for imaging objects with fast dynamics, since high brightness is favorable in short acquisition time required by the rapid movement. On the contrary, SRhB with prolonged on time and photostability is more suitable for imaging objects with relative slow movements, as long acquisition time could be utilized to collect sufficient photons in regard to the prolonged on time.

## Live-cell Super-resolution Imaging

Following the predictions, we next sought to experimentally investigate the super-resolution imaging performance of SRhB and STMR on various targets in living cells with the aid of Halo-tag technology. Four typical cellular organelles were selected: endoplasmic reticulum (ER)^47–50^ and mitochondria,^51–53^ which are highly plastic and exhibit rapid movement; nucleus and microtubule, which are relatively static during imaging time (< 3 min). To label these organelles and avoid expression disparity, single-cell clone was deployed to construct three cell lines (HeLa or U2OS) stably expressing Halo-Sec61β (endoplasmic reticulum, ER), Halo-TOMM20 (mitochondrial outer membrane) or Halo-H2B (nucleus) fusion proteins. The preparation of Halo-β-tubulin (microtubule) stably expressing cell line was failed in our study probably owing to the division disordering of transfected cells, and transient expressing cells were utilized instead. Both STMR and SRhB were specifically labeled to target fusion proteins (Figure S3) in living cells, confirming their membrane permeability. During super-resolution imaging acquisition, both fluorophores solidly demonstrated large number of spontaneous blinking events, in accordance with their self-blinking characteristics. Through localizing the event positions, the morphology of ER (Figure 3a, b), mitochondria (Figure 3c, d), the distribution of H2B proteins in nucleus (Figure 3e, f) and the twisted structure of microtubules (Figure 3g, h) were successfully reconstructed. Moreover, the magnified views of reconstructions reveal the grid structure of ER (Figure 3a, b insets) and rod structure of mitochondria (Figure 3c, d insets), with definite clarity enhancement compared to the conventional imaging (the sharp contrast was also validated for imaging H2B proteins and microtubules in Figure S3).

**Figure 3.**
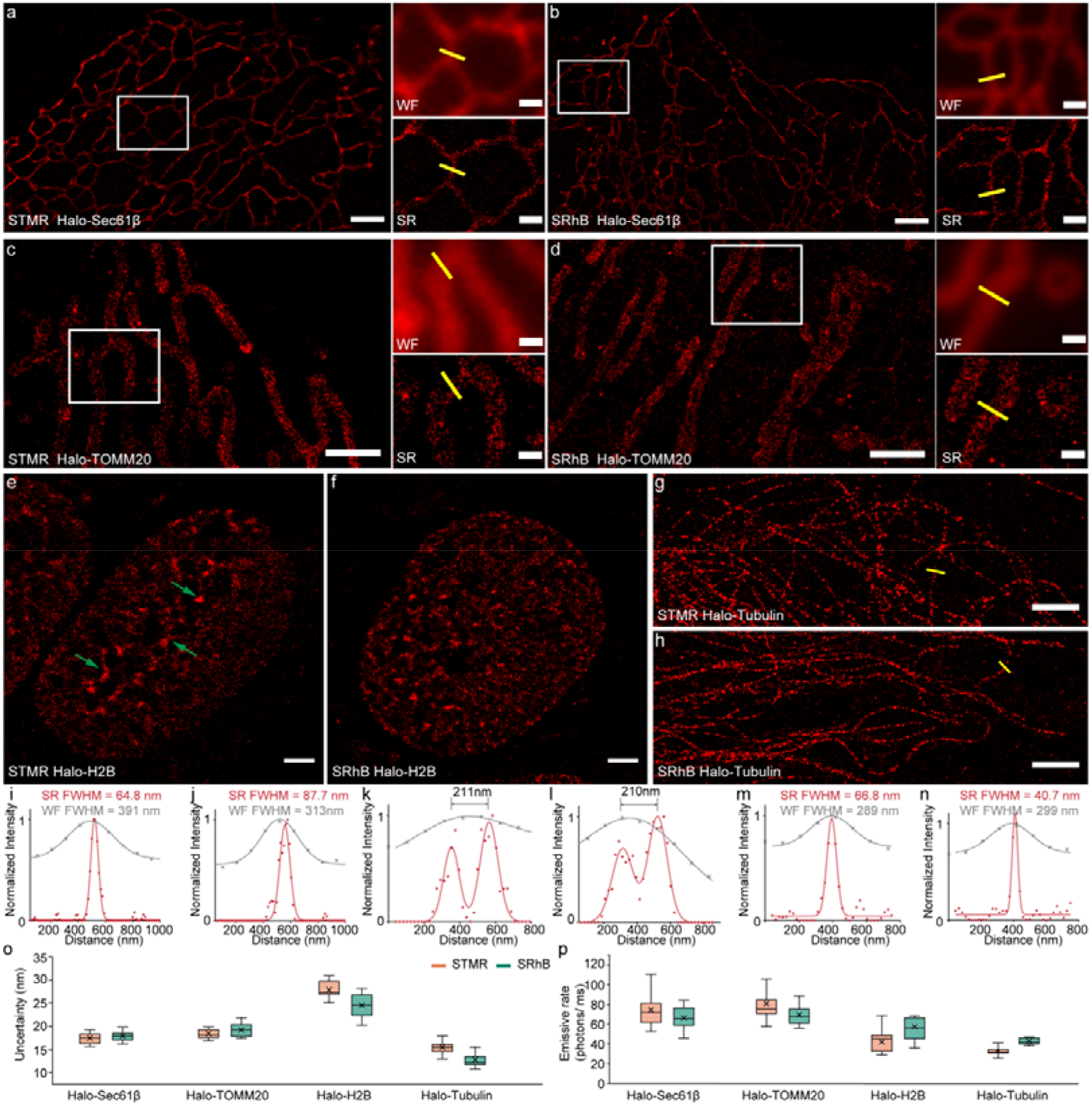
Comparison on the super-resolution imaging of endoplasmic reticulum (a-b), mitochondria (c-d), H2B proteins (e-f) and microtubules (g-h) in living cells stained with STMR and SRhB. Insets on right panel shows the conventional image (WF) and the super-resolution image (SR) of a magnified view at the highlighted regions in (a-d). (i-n) Intensity profiles of endoplasmic reticulum, mitochondria and microtubules indicated with yellow lines in super-resolution image (a, b, c, d, g and h). The box plots of localization uncertainties (o) and emissive rate (p) for STMR and SRhB on reconstructed cellular organelles (cell number > 15). Scale bars: 2 μm (a-f); 1 μm (g, h); 0. 5 μm (Insets).

To further compare the imaging capabilities of the two fluorophores, the reconstruction is quantitatively evaluated and compared. For imaging ER and mitochondria with rapid dynamics, STMR is superior than SRhB with an enhanced resolution (Figure S4-S5). Typically, in super-resolution imaging ER, the width of ER tubules is estimated at 60 nm for STMR (Figure 3i), narrower than that of SRhB (80 nm; Figure 3j); for imaging mitochondria, STMR precisely separated the mitochondrial outer membranes distanced at 211 nm, while that of SRhB is barely vague (Figure 3k-l). On the other side, the results support a suitability of SRhB for imaging H2B proteins and microtubules compared to STMR (Figure S6-S7). Although the same cell line (from single-cell clone) was used, the H2B proteins shown different distribution patterns in the reconstructions from STMR and SRhB. The abnormal clusters (marked by green arrows in Figure 3e) of H2B proteins in the reconstruction indicate a poor imaging quality for STMR. For imaging microtubules, the width of tubular fiber is narrower for SRhB (40.7 nm; Figure 3n) than that of STMR (66.8 nm; Figure 3m). Furthermore, the resolution disparities are reflected by the localization errors for the two fluorophores (Figure 3o; ER, 17.5 *vs* 18; mitochondria, 18.3 *vs* 19.3; H2B, 27.5 *vs* 24.8; microtubule, 15.6 *vs* 12.3; unit is nm, order in STMR *vs* SRhB). Although the self-blinking of STMR and SRhB provides sufficient localizations to plot the target structures, the reconstructions from the two demonstrate varied resolution levels. Next, this disparity is analyzed in regard to the imaging requirements and the single-molecule characteristics.

In in-vitro analysis, single-molecule events of STMR exhibit a photon burst at high brightness, but their on-time and photostability are inferior to those of SRhB. Hence, STMR is potentially suitable for imaging at a short exposure time (< 10 ms), while SRhB is ready for providing consistent emission at a long exposure time (20-100 ms), which could increase the brightness of single-molecule events. On the other hand, every targets in living cells has its own requirements for imaging methods. The ER and mitochondria are rapid transforming organelles, and their imaging required a high temporal resolution (< 5 s; 1000 frames at 200 Hz). Such high temporal resolution would be benefited from a short exposure time of single-molecule events, as the reconstructions are incomplete at a long exposure time (125 frames at 25 Hz; Figure S8). The movements of the structural H2B proteins and microtubule are negligible in long imaging time (1-2 min; 1500-3000 frames at 25 Hz). If short exposure time (100 Hz) is applied, the insufficient collection of photons for single-molecule events could result in large localizing errors, blurring the reconstruction (Figure S9). Therefore, it is unambiguously inferred that STMR is more suitable for ER and mitochondrial imaging than SRhB, while SRhB is more suitable for nucleus and microtubule imaging. This prediction is experimentally confirmed with emissive rates of single-molecule events during imaging (Figure 3p; ER, 74 *vs* 63; mitochondria, 66 *vs* 59; H2B, 38 *vs* 48; microtubule, 23 *vs* 33; unit is photons/ms, order in STMR *vs* SRhB). These differences solidly constitute the resolution disparity of reconstruction for the two fluorophores. Through the distinct imaging results from STMR and SRhB under identical conditions, it demonstrates that subtle structural differences, even as small as a methyl group, could raise a butterfly effect to influence where a fluorophore is best suited for imaging.

## Resolving Mitochondrial Transforms Under Apoptosis and Ferroptosis

Mitochondria adaptively transform morphologies in various programmed cell death pathways (apoptosis, ferroptosis and etc.), helping defensing the hosts from pathogens, cancer and neuro degenerative disease.^54–56^ Yet, the ultra-details of mitochondria in the process of cell programmed death are lost in previous mitochondrial studies due to diffraction limit, which constitutes a barrier for further understanding the interconnectivity and molecular machinery of various pathways.

Herein, mitochondrial morphological under apoptosis and ferroptosis were successfully monitored with super-resolution imaging for the first time (Figure 4). To implement this imaging, STMR instead of SRhB is selected on the basis of the high brightness and imaging suitability, and U2OS cell stably expressing Halo-TOMM20 proteins are recruited. In apoptosis induced by carbonyl cyanide m-chlorophenylhydrazine (CCCP), the tubular morphology of normal mitochondria (Figure 4a) immediately break and shorten (Figure 4b) in just 0.5 h. At 2 h, almost all mitochondria (>80%) turn to sphere fragments showing round projection (Figure 4c) on the reconstructions. Magnified views on a round mitochondrion (Figure 4c insets) further reveal the clarity enhancement from reconstruction to the conventional images. In contrast to the apoptosis process, mitochondria transforms slowly in the ferroptosis pathway stimulated by erastin. None significant change on mitochondrial morphology is observed at 2 h. Mitochondria begin to break and shorten until 4 h of stimulation (Figure 4d), and finally shrunk to diversified mitochondrial fragments with no more significant changes after 8 h (Figure 4e, f). One of the large mitochondrial fragments is magnified in Figure 4f insets, with bending structures. The formed round fragments in ferroptosis shown smaller-sized area projection, and many fragments (43%) show irregular shapes compared to those obtained in apoptosis. In accordance with above observations, the quantitative statistics of mitochondrial length (Figure 4g) and shape (Figure 4h) confirmed the morphological transforms of mitochondria in both pathways. Apoptosis show fast speed in forming the fragments, which is credited to the associating outer mitochondrial membrane permeabilization. On the other side, the round-shaped mitochondrial fragments formed in ferroptosis are a surprise discovery, which show morphological similarity to those formed in apoptosis, and might suggest the interconnectivity of two pathways. Yet, the diversified structures of fragments indicate the ferroptosis is rather complex with the involvements of other undefined pathways.

**Figure 4.**
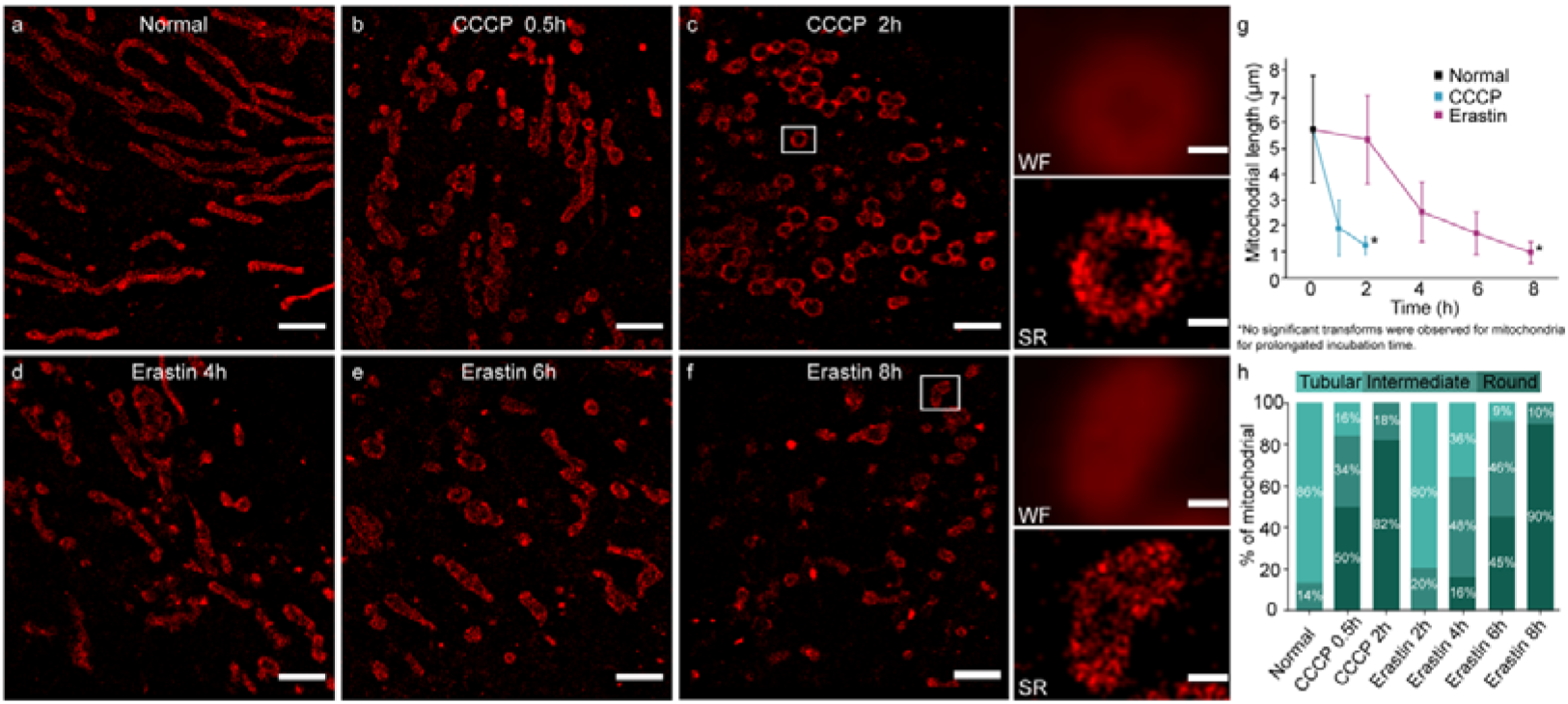
(a-f) The morphological change of mitochondria during the apoptosis and ferroptosis in U2OS cells expressing Halo-TOMM20 treated with CCCP (10 μM)/ erastin (10 μM) for the indicated times. Insets on right panel shows the conventional image (WF) and the super-resolution image (SR) of a magnified view at the highlighted regions in (c, f). The box plots of mitochondrial length (g) and quantitative analysis of mitochondrial morphology (h) during the apoptosis and ferroptosis. Scale bars: 2 μm (a-f); 0.25 μm (Insets).

## Conclusions

In conclusion, to remind the researchers that a smallest structural change in fluorophores may raise a great difference in single-molecule localization super-resolution imaging, we designed a pair of homologous sulforhodamine dyes, i.e. SRhB and STMR with ethyl and methyl substituents, respectively. Due to their almost identical structures, SRhB and STMR exhibited proximate absoprtion, fluorescent spectra and spirolactam-zwitterion equilibrium coefficients (p*K*_cycling_). However, they differed in their single-molecule fluorescence characteristics. On one side, both SRhB and STMR demonstrated fast recruiting rates and large numbers of switching, suggesting their self-blinking functionality for SMLM imaging. On the other side, their switching events shown unique properties: single-molecule events of STMR showed a higher single-molecule brightness, while those of SRhB presented prolonged on-time and photostability. This uniqueness suggests individual suitability of STMR and SRhB for SMLM imaging, which should be wisely selected to match the diversified targets with various dynamics in living cells. To provide experimental demonstration, through the Halo protein-tag technology, SMLM imaging capability of STMR and SRhB were compared under identical microenvironments on context of labeling endoplasmic reticulum, outer mitochondrial membranes, H2B proteins and microtubules in live cells. For highly plastic and fast-moving subcellular organelles, i.e. endoplasmic reticulum and mitochondria, STMR exhibited superior imaging quality than SRhB, since the former has a higher emissive rate. Correspondingly, for organelles, i.e. nucleus and microtubules, relatively kept static during imaging time, the reconstruction could benefited from prolonged on time of SRhB by deploying long accumulating time for single-molecule events, which assist in providing enhanced clarity and integrity. With these inspiring knowledges, STMR instead of SRhB is employed to first super-resolve the morphological changes of the mitochondrial outer membrane during apoptosis and ferroptosis. Our study would remind scientists that even a subtle methyl group difference in structure may cause a butterfly effect, leading to a large variance in their performance for single-molecule super-resolution imaging. We foresee that our results would provide guidance for the future design of fluorophores for localization-based imaging, and entail the rational selection of fluorophores for single-molecule imaging in different structural environments in the future.

## ASSOCIATED CONTENT

### Supporting Information

The Supporting Information is available free of charge on the xxxxx at DOI: xxxxxx

## AUTHOR INFORMATION

### Notes

Any additional relevant notes should be placed here.

## ACKNOWLEDGMENT

This work was supported by the National Natural Science Foundation of China (Nos. 21804016, 22004011, and 22174009), the China Postdoctoral Science Foundation (Nos. BX20200073 and 2020M670754), the Fundamental Research Funds for the Central Universities (Nos. DUT22LAB608, DUT20JC39 and DUT21YG126), and the Dalian Science and Technology Innovation Fund (No. 2020JJ25CY014).

## REFERENCES

1. Hell, S. W.; Wichmann, J., Breaking the diffraction resolution limit by stimulated emission: stimulated-emission-depletion fluorescence microscopy. Optics letters 1994, 19 (11), 780–2.

2. Sahl, S. J.; Hell, S. W.; Jakobs, S., Fluorescence nanoscopy in cell biology. Nat. Rev. Mol. Cell Biol. 2017, 18 (11), 685–701.

3. Huang, B.; Babcock, H.; Zhuang, X., Breaking the diffraction barrier: super-resolution imaging of cells. Cell 2010, 143 (7), 1047–58.

4. Fornasiero, E. F.; Opazo, F., Super-resolution imaging for cell biologists: concepts, applications, current challenges and developments. BioEssays : news and reviews in molecular, cellular and developmental biology 2015, 37 (4), 436–51.

5. van de Linde, S.; Heilemann, M.; Sauer, M., Live-cell super-resolution imaging with synthetic fluorophores. Annu. Rev. Phys. Chem. 2012, 63, 519–540.

6. Ha, T.; Tinnefeld, P., Photophysics of Fluorescent Probes for Single-Molecule Biophysics and Super-Resolution Imaging. Annu. Rev. Phys. Chem. 2012, 63 (1), 595–617.

7. Minoshima, M.; Kikuchi, K., Photostable and photoswitching fluorescent dyes for super-resolution imaging. JBIC Journal of Biological Inorganic Chemistry 2017, 22 (5), 639–652.

8. Fernández-Suárez, M.; Ting, A. Y., Fluorescent probes for super-resolution imaging in living cells. Nat. Rev. Mol. Cell Biol. 2008, 9 (12), 929–943.

9. Vogelsang, J.; Steinhauer, C.; Forthmann, C.; Stein, I. H.; Person-Skegro, B.; Cordes, T.; Tinnefeld, P., Make them Blink: Probes for Super-Resolution Microscopy. ChemPhysChem 2010, 11 (12), 2475–2490.

10. Li, H.; Vaughan, J. C., Switchable Fluorophores for Single-Molecule Localization Microscopy. Chem. Rev. 2018, 118 (18), 9412–9454.

11. Dempsey, G. T.; Vaughan, J. C.; Chen, K. H.; Bates, M.; Zhuang, X., Evaluation of fluorophores for optimal performance in localization-based super-resolution imaging. Nat. Methods 2011, 8 (12), 1027–1036.

12. Jradi, F. M.; Lavis, L. D., Chemistry of Photosensitive Fluorophores for Single-Molecule Localization Microscopy. ACS Chemical Biology 2019, 14 (6), 1077–1090.

13. Lincoln, R.; Bossi, M. L.; Remmel, M.; D’Este, E.; Butkevich, A. N.; Hell, S. W., A general design of caging-group-free photoactivatable fluorophores for live-cell nanoscopy. Nat. Chem. 2022, 14 (9), 1013–1020.

14. Qi, Q.; Chi, W.; Li, Y.; Qiao, Q.; Chen, J.; Miao, L.; Zhang, Y.; Li, J.; Ji, W.; Xu, T.; Liu, X.; Yoon, J.; Xu, Z., A H-bond strategy to develop acid-resistant photoswitchable rhodamine spirolactams for super-resolution single-molecule localization microscopy. Chemical Science 2019, 10 (18), 4914–4922.

15. Butkevich, A. N.; Bossi, M. L.; Lukinavičius, G.; Hell, S. W., Triarylmethane Fluorophores Resistant to Oxidative Photobluing. J. Am. Chem. Soc. 2019, 141 (2), 981–989.

16. Tyson, J.; Hu, K.; Zheng, S.; Kidd, P.; Dadina, N.; Chu, L.; Toomre, D.; Bewersdorf, J.; Schepartz, A., Extremely Bright, Near-IR Emitting Spontaneously Blinking Fluorophores Enable Ratiometric Multicolor Nanoscopy in Live Cells. ACS Central Science. 2021, 7 (8), 1419–1426.

17. Zheng, Y.; Ye, Z.; Liu, Z.; Yang, W.; Zhang, X.; Yang, Y.; Xiao, Y., Nitroso-Caged Rhodamine: A Superior Green Light-Activatable Fluorophore for Single-Molecule Localization Super-Resolution Imaging. Anal. Chem. 2021, 93 (22), 7833–7842.

18. Ye, Z.; Zheng, Y.; Peng, X.; Xiao, Y., Surpassing the Background Barrier for Multidimensional Single-Molecule Localization Super-Resolution Imaging: A Case of Lysosome-Exclusively Turn-on Probe. Anal. Chem. 2022, 94 (22), 7990–7995.

19. Ye, Z.; Yu, H.; Yang, W.; Zheng, Y.; Li, N.; Bian, H.; Wang, Z.; Liu, Q.; Song, Y.; Zhang, M.; Xiao, Y., Strategy to Lengthen the On-Time of Photochromic Rhodamine Spirolactam for Super-resolution Photoactivated Localization Microscopy. J. Am. Chem. Soc. 2019, 141 (16), 6527–6536.

20. Zheng, Q.; Ayala, A. X.; Chung, I.; Weigel, A. V.; Ranjan, A.; Falco, N.; Grimm, J. B.; Tkachuk, A. N.; Wu, C.; Lippincott-Schwartz, J.; Singer, R. H.; Lavis, L. D., Rational Design of Fluorogenic and Spontaneously Blinking Labels for Super-Resolution Imaging. ACS Central Science. 2019, 5 (9), 1602–1613.

21. Grimm, F.; Rehman, J.; Stoldt, S.; Khan, T. A.; Schlötel, J. G.; Nizamov, S.; John, M.; Belov, V. N.; Hell, S. W., Rhodamines with a Chloronicotinic Acid Fragment for Live Cell Superresolution STED Microscopy**. Chem. - Eur. J. 2021, 27 (19), 6070–6076.

22. Montenegro, H.; Di Paolo, M.; Capdevila, D.; Aramendía, P. F.; Bossi, M. L., The mechanism of the photochromic transformation of spirorhodamines. Photochemical & Photobiological Sciences 2012, 11 (6), 1081–1086.

23. Uno, S.-n.; Kamiya, M.; Yoshihara, T.; Sugawara, K.; Okabe, K.; Tarhan, M. C.; Fujita, H.; Funatsu, T.; Okada, Y.; Tobita, S.; Urano, Y., A spontaneously blinking fluorophore based on intramolecular spirocyclization for live-cell super-resolution imaging. Nat. Chem. 2014, 6 (8), 681–689.

24. Ye, Z.; Yang, W.; Wang, C.; Zheng, Y.; Chi, W.; Liu, X.; Huang, Z.; Li, X.; Xiao, Y., Quaternary Piperazine-Substituted Rhodamines with Enhanced Brightness for Super-Resolution Imaging. J. Am. Chem. Soc. 2019, 141 (37), 14491–14495.

25. Wang, L.; Tran, M.; D’Este, E.; Roberti, J.; Koch, B.; Xue, L.; Johnsson, K., A general strategy to develop cell permeable and fluorogenic probes for multicolour nanoscopy. Nat. Chem. 2020, 12 (2), 165–172.

26. Wang, B.; Xiong, M.; Susanto, J.; Li, X.; Leung, W.-Y.; Xu, K., Transforming Rhodamine Dyes for (d)STORM Super-Resolution Microscopy via 1,3-Disubstituted Imidazolium Substitution. Angew. Chem., Int. Ed. 2022, 61 (9), e202113612.

27. Beija, M.; Afonso, C. A. M.; Martinho, J. M. G., Synthesis and applications of Rhodamine derivatives as fluorescent probes. Chem. Soc. Rev. 2009, 38 (8), 2410–2433.

28. Klein, U. K. A.; Hafner, F. W., A new dual fluorescence with rhodamine B lactone. Chemical Physics Letters 1976, 43 (1), 141–145.

29. Ravdin, P.; Axelrod, D., Fluorescent tetramethyl rhodamine derivatives of α-bungarotoxin: Preparation, separation, and characterization. Analytical Biochemistry 1977, 80 (2), 585–592.

30. Vult von Steyern, F.; Josefsson, J. O.; Tågerud, S., Rhodamine B, a fluorescent probe for acidic organelles in denervated skeletal muscle. Journal of Histochemistry & Cytochemistry 1996, 44 (3), 267–274.

31. Savarese, M.; Aliberti, A.; De Santo, I.; Battista, E.; Causa, F.; Netti, P. A.; Rega, N., Fluorescence Lifetimes and Quantum Yields of Rhodamine Derivatives: New Insights from Theory and Experiment. The Journal of Physical Chemistry A 2012, 116 (28), 7491–7497.

32. Belov, V. N.; Bossi, M. L., Photoswitching Emission with Rhodamine Spiroamides for Super-resolution Fluorescence nanoscopies. Israel Journal of Chemistry 2013, 53 (5), 267–279.

33. Liu, Z.; Zheng, Y.; Xie, T.; Chen, Z.; Huang, Z.; Ye, Z.; Xiao, Y., Clickable rhodamine spirolactam based spontaneously blinking probe for super-resolution imaging. Chinese Chemical Letters 2021, 32 (12), 3862–3864.

34. Xu, Y.; Xu, R.; Wang, Z.; Zhou, Y.; Shen, Q.; Ji, W.; Dang, D.; Meng, L.; Tang, B. Z., Recent advances in luminescent materials for super-resolution imaging via stimulated emission depletion nanoscopy. Chem. Soc. Rev. 2021, 50 (1), 667–690.

35. Bates, M.; Huang, B.; Zhuang, X., Super-resolution microscopy by nanoscale localization of photo-switchable fluorescent probes. Current Opinion in Chemical Biology 2008, 12 (5), 505–514.

36. Banala, S.; Maurel, D.; Manley, S.; Johnsson, K., A Caged, Localizable Rhodamine Derivative for Superresolution Microscopy. ACS Chemical Biology 2012, 7 (2), 289–293.

37. Grimm, J. B.; English, B. P.; Choi, H.; Muthusamy, A. K.; Mehl, B. P.; Dong, P.; Brown, T. A.; Lippincott-Schwartz, J.; Liu, Z.; Lionnet, T.; Lavis, L. D., Bright photoactivatable fluorophores for single-molecule imaging. Nat. Methods 2016, 13 (12), 985–988.

38. He, H.; Ye, Z.; Xiao, Y.; Yang, W.; Qian, X.; Yang, Y., Super-Resolution Monitoring of Mitochondrial Dynamics upon Time-Gated Photo-Triggered Release of Nitric Oxide. Anal. Chem. 2018, 90 (3), 2164–2169.

39. Werther, P.; Yserentant, K.; Braun, F.; Kaltwasser, N.; Popp, C.; Baalmann, M.; Herten, D.-P.; Wombacher, R., Live-Cell Localization Microscopy with a Fluorogenic and Self-Blinking Tetrazine Probe. Angew. Chem., Int. Ed. 2020, 59 (2), 804–810.

40. Takakura, H.; Zhang, Y.; Erdmann, R. S.; Thompson, A. D.; Lin, Y.; McNellis, B.; Rivera-Molina, F.; Uno, S.-n.; Kamiya, M.; Urano, Y.; Rothman, J. E.; Bewersdorf, J.; Schepartz, A.; Toomre, D., Long time-lapse nanoscopy with spontaneously blinking membrane probes. Nat. Biotechnol. 2017, 35 (8), 773–780.

41. Gerasimaitė, R. t.; Bucevičius, J.; Kiszka, K. A.; Schnorrenberg, S.; Kostiuk, G.; Koenen, T.; Lukinavičius, G., Blinking Fluorescent Probes for Tubulin Nanoscopy in Living and Fixed Cells. ACS Chemical Biology 2021, 16 (11), 2130–2136.

42. Zheng, Y.; Ye, Z.; Xiao, Y., Turn-on Rate Determines the Blinking Propensity of Rhodamine Fluorophores for Super-Resolution Imaging. bioRxiv 2022, 2022.10.17.512490.

43. Los, G. V.; Encell, L. P.; McDougall, M. G.; Hartzell, D. D.; Karassina, N.; Zimprich, C.; Wood, M. G.; Learish, R.; Ohana, R. F.; Urh, M.; Simpson, D.; Mendez, J.; Zimmerman, K.; Otto, P.; Vidugiris, G.; Zhu, J.; Darzins, A.; Klaubert, D. H.; Bulleit, R. F.; Wood, K. V., HaloTag: A Novel Protein Labeling Technology for Cell Imaging and Protein Analysis. ACS Chemical Biology 2008, 3 (6), 373–382.

44. Miller, L. W.; Cai, Y.; Sheetz, M. P.; Cornish, V. W., In vivo protein labeling with trimethoprim conjugates: a flexible chemical tag. Nat. Methods 2005, 2 (4), 255–257.

45. Griffin, B. A.; Adams, S. R.; Tsien, R. Y., Specific Covalent Labeling of Recombinant Protein Molecules Inside Live Cells. Science 1998, 281 (5374), 269–272.

46. Jing, C.; Cornish, V. W., Chemical Tags for Labeling Proteins Inside Living Cells. Accounts of Chemical Research. 2011, 44 (9), 784–792.

47. Nixon-Abell, J.; Obara, C. J.; Weigel, A. V.; Li, D.; Legant, W. R.; Xu, C. S.; Pasolli, H. A.; Harvey, K.; Hess, H. F.; Betzig, E.; Blackstone, C.; Lippincott-Schwartz, J., Increased spatiotemporal resolution reveals highly dynamic dense tubular matrices in the peripheral ER. Science 2016, 354 (6311), aaf3928.

48. Schwarz, D. S.; Blower, M. D., The endoplasmic reticulum: structure, function and response to cellular signaling. Cellular and Molecular Life Sciences 2016, 73 (1), 79–94.

49. Lee, C.; Chen, L. B., Dynamic behavior of endoplasmic reticulum in living cells. Cell 1988, 54 (1), 37–46.

50. Perkins, H. T.; Allan, V., Intertwined and Finely Balanced: Endoplasmic Reticulum Morphology, Dynamics, Function, and Diseases. Cells 2021, 10 (9), 2341.

51. Fehrenbacher, K. L.; Yang, H.-C.; Gay, A. C.; Huckaba, T. M.; Pon, L. A., Live Cell Imaging of Mitochondrial Movement along Actin Cables in Budding Yeast. Current Biology 2004, 14 (22), 1996–2004.

52. Mitra, K.; Lippincott-Schwartz, J., Analysis of Mitochondrial Dynamics and Functions Using Imaging Approaches. Current Protocols in Cell Biology 2010, 46 (1), 4.25.1–4.25.21.

53. Kuznetsov, A. V.; Hermann, M.; Saks, V.; Hengster, P.; Margreiter, R., The cell-type specificity of mitochondrial dynamics. The International Journal of Biochemistry & Cell Biology 2009, 41 (10), 1928–1939.

54. Wai, T.; Langer, T., Mitochondrial Dynamics and Metabolic Regulation. Trends in Endocrinology & Metabolism 2016, 27 (2), 105–117.

55. Youle, R. J.; van der Bliek, A. M., Mitochondrial Fission, Fusion, and Stress. Science 2012, 337 (6098), 1062–1065.

56. Friedman, J. R.; Nunnari, J., Mitochondrial form and function. Nature 2014, 505 (7483), 335–343.

