## supporting information for "Subtle Structural Translation Remarkably Modulates the Super-Resolution Imaging of Self-blinking Rhodamines"

### 1 Synthetic method

#### 1.1 General

All regents, e.g. MeOH, DMF, acetonitrile, triethylamine, etc., were purchased from commercial suppliers and used as received. Column chromatography was performed with silica gel (200-300 mesh and 300-400 mesh).

$^{1}H$ NMR and $^{13}C$ NMR were measured on Bruker Avance II 400, Bruker Avance III 500 and Varian MERCURY 400 spectrometers. Mass spectra and high-resolution mass spectra were recorded on HP 1100 LC-MSD, gas chromatography/TOF Mass, thermo Scientific LTQ Orbitrap XL, Waters Synapt G2-Si HDMS and UPLC/Q-TOF Mass spectrometers.

#### 1.2 Synthesis of SRhB-COOH

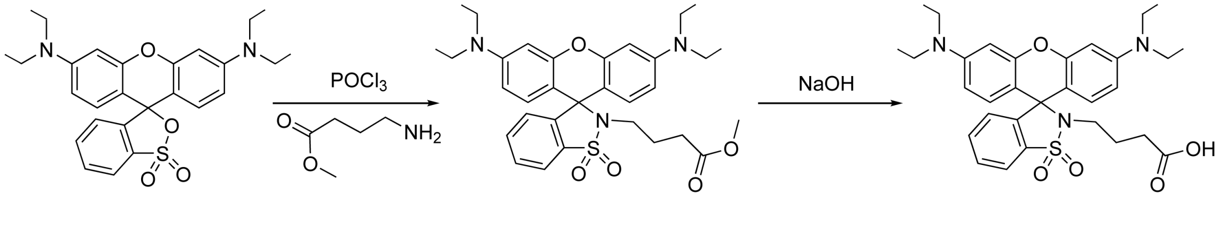

SRhB-COOMe SRhB-COOH

SRhB-COOH: Sulforhodamine B (100 mg, 0.209 mmol) was dissolved in dry 1,2-dichloroethane (5 mL) and stirred vigorously. Phosphorus oxychloride (25 μL, 0.21 mmol) was added at room temperature for 5 min. Then the solution was refluxed for 2h. The obtained mixture was cooled for the next step. The mixture of crude acid chloride was added to a DCM/MeCN (5mL/5 mL) mixture solution of 4-amino-butyricacimethylester (29 mg, 0.25 mmol) and triethylamine at ice bath. After stirring 2h, the crude product was purified through silica gelcolumn chromatography with a mixture of petroleum ether and ethylene acetate (5:1), v/v) as eluent. SRhB-COOMe was obtained as a colorless powder (90 mg, yield 74%). SRhB-COOMe (90mg, 0.16 mmol) and NaOH (33mg, 0.81 mmol) was dissolved in methane/H2O (8mL/2mL) and refluxed 3h. After completion of the reaction (monitored via thin-layer chromatography), the solution was evaporated in vacuo. SRhB-COOH was obtained through silica gel column chromatography with a mixture of dichloromethane and methane (10:1, v/v) as eluent. The collected solution was dried in vacuo to afford STMR-COOH (80 mg, yield 91%). 1H NMR (400 MHz, DMSO-d6) δ 8.25 – 7.87 (m, 1H), 7.78 – 7.36 (m, 2H), 7.13 – 6.85 (m, 1H), 6.66 (d, J = 8.9 Hz, 2H), 6.54 – 6.40 (m, 2H), 6.38 – 6.31 (m, 2H), 3.33 (q, J = 7.8, 6.9 Hz, 8H), 2.82 (t, J = 6.9 Hz, 2H), 2.00 (t, J = 7.4 Hz, 2H), 1.45 (p, J = 7.4, 6.6 Hz, 2H), 1.09 (t, J = 6.8 Hz, 12H). $^{13}C$ NMR (101 MHz, DMSO) δ 174.20, 152.70, 148.98, 145.67, 134.41, 133.07, 129.96, 129.68, 126.65, 120.53, 108.82, 106.57, 97.33, 66.14, 55.38, 44.14, 31.20, 24.09, 12.86. HRMS (ESI) *m*/*z* called for $C_{31}H_{38}N_3O_5S$ $[M+H]^+$: 564.2527; found: 564.2526 ($z$ = 1).

1.3 Synthesis of SRhB

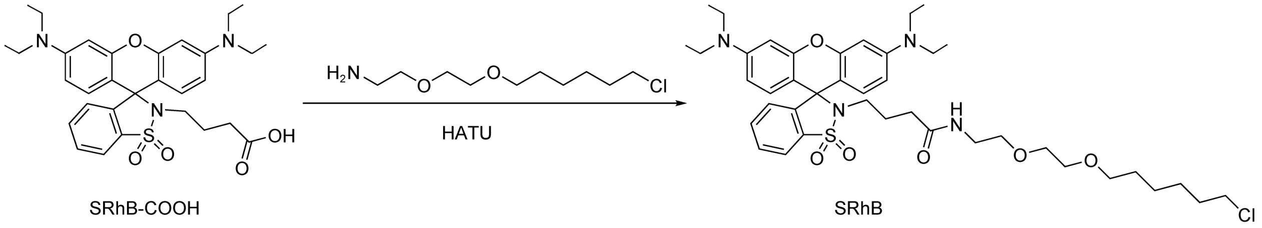

SRhB: SRhB-COOH (55mg, 0.098 mmol), Halo-$NH_2$ (27mg, 0.12 mmol), and HATU (45mg, 0.12 mmol) were dissolved in dry N,N-dimethylformamide (2 mL) in the presence of N,N-disopropylethylamine (20 μL, 0.12 mmol). The mixture was stirred at room temperature for 1h. The reaction mixture was further washed with brine, dried over $Na_2SO_4$, filtered and evaporated. The crude product was purified through silica gel column chromatography (dichloromethane: methane=30:1). SRhB (light pink powder, 33mg, yield 44%) was obtained. $^1$H NMR (400 MHz, DMSO-$d_6$) δ 8.02 – 7.89 (m, 1H), 7.74 – 7.41 (m, 2H), 6.95 (dd, *J* = 5.3, 3.7 Hz, 1H), 6.69 (d, *J* = 8.9 Hz, 2H), 6.44 (dd, *J* = 9.0, 2.6 Hz, 2H), 6.35 (d, *J* = 2.5 Hz, 2H), 3.61 (t, *J* = 6.6 Hz, 2H), 3.44 (s, 4H), 3.40 – 3.30 (m, 8H), 3.29 (d, *J* = 5.9 Hz, 4H), 3.06 (q, *J* = 6.0 Hz, 2H), 2.85 – 2.71 (m, 2H), 1.88 (t, *J* = 7.4 Hz, 2H), 1.69 (dt, *J* = 14.4, 6.6 Hz, 2H), 1.48 (dq, *J* = 13.7, 6.8 Hz, 4H), 1.41 – 1.32 (m, 2H), 1.31 – 1.24 (m, 2H), 1.09 (t, *J* = 7.0 Hz, 12H) 13C NMR (126 MHz, DMSO) δ 171.16, 152.17, 148.45, 145.29, 133.87, 132.43, 129.39, 129.15, 126.07, 120.01, 108.27, 106.15, 96.84, 70.13, 69.51, 69.34, 68.97, 65.63, 54.87, 45.30, 43.61, 38.33, 32.59, 31.97, 29.01, 26.06, 24.87, 24.69, 12.37. HRMS (ESI) *m*/*z* called for $C_{41}H_{58}ClN_4O_6S$ $[M+H]^+$: 769.3766; found: 769.3769 (*z* = 1).

#### 2 Spectroscopic study

Absorption spectra were recorded on Agilent 8453 UV-visible spectrophotometer and fluorescence spectra were recorded on Agilent Cary Eclipse Fluorescence spectrophotometer.

2.1 Method

2.1.1 Calculation of $pK_{cycling}$ value.

STMR and SRhB were prepared as 2 μM solution for measurements. Fluorescence spectra was measured in PBS (10 mM) containing 10% EtOH at various pH values.

2.1.2 Measurement of molecular kinetics in vitro.

SRhB were prepared as 2 μM solution in MilliQ water containing 30% EtOH for measurements. A mixed buffer was rapidly changed from an acidic environment around pH 4 to an alkaline environment at pH 10 to detect the change of the fluorescence intensity of SRhB with time. Another mixed buffer was rapidly changed from pH 10 to pH 4 in an acidic environment to detect the fluorescence intensity of SRhB as a function of time.

2.2 Spectral Analysis

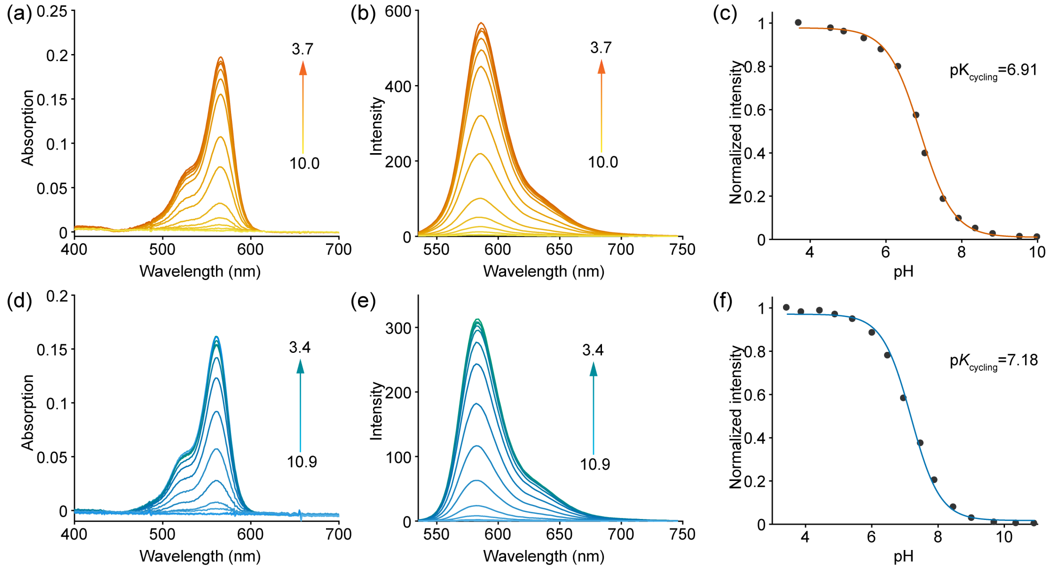

**Figure S1**. Protonation p$K_{cycling}$ measurements for SRhB (top panel, a-c) and STMR (bottom panel, d-f) in PBS/EtOH (v/v = 9: 1). The absorption spectra changes (a and d) and the corresponding fluorescent spectra changes (b and e) and integrated emission intensity (c and f) as a function of pH.

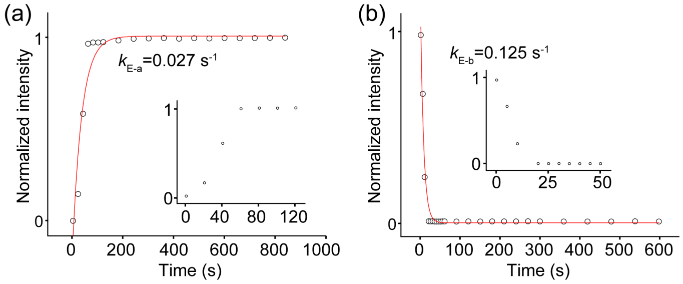

**Figure S2**. Ensemble kinetic study of spirocyclization equilibrium after acid or basic perturbation. Peak emission intensities (582 nm) are plotted as a function of time, showing the rates reflecting equilibrium shifts to ring-opening ($k_{E-a}$, a) and ring-closing states ($k_{E-b}$, b).

### 3 Single-Molecule study

#### 3.1 Microscopy

Single-molecule and Super-resolution imaging were studied with a total internal reflection fluorescence microscope (TIRFM) built on an Olympus IX71 inverted microscope as described earlier.[1, 2] The laser light was focused on the back focal plane of a x 100 objective (UAPON 100XOTIRF; 1.49 numerical aperture). An EMCCD camera (iXon DU-897U) was implemented for data acquisition.

#### 3.2 Protein Labeling

SRhB was labeled to Halo protein in PBS (pH=7.4), then the resulting mixture was incubated at 37 ℃ for 2h. The remaining unbounded fluorophores were removed by protein desalting spin columns (filled with Sephadex G-25 resins).

#### 3.3 Single-Molecule Imaging

The protein solution labeled with SRhB were freshly diluted in PBS at a low concentration to minimize the overlapping between different molecules and transferred to the surface of clean coverslips. The adhesion of proteins to the surface proceed for 30 s and the unbounded proteins were washed out through three times rinses of PBS. The single-molecule fluorescence was recorded with the microscopy described above in conventional mode. The pixel size was 160 nm/pixel and 6000 raw frames were acquired at 10 ms exposure time under different laser power irradiation. At least five measurements were performed for each irradiation condition.

3.4 Single-Molecule Analysis

Raw data were automatically processed with a home-written MatLab software as described in our previous report.[2-4] Briefly, the molecular candidates were identified by denoising. The spatially approximate molecules were removed during this process. Then the fluorescent trajectories of these molecules were extracted and fitted to a hidden Markov model with a Gaussian distribution as the observation probability distribution.

*Brightness.* Single-molecule brightness in this manuscript was determined as the photon counts from a single molecule during the acquisition time of single frame (10 ms).

*On time.* The time duration of a bright state.

*Switch number.* The counted times of bright-to dark transitions of a molecule.

*Recruiting rate.* The rate at which a molecule transitions from the initial dark state to the bright state. (Rhodamine molecules in initial dark state probably exist in their ring-closed form at ground state, as those molecules have not been excited and shown none fluorescence from the beginning.)

### 4 Super-resolution Imaging

4.1 cell culture

HeLa cells were purchased from the Cell Bank of Type Culture of Chinese Academy of Sciences. Vero and U2OS cells were kindly provided by Procell Life Science&Technology Co.,Ltd. HeLa and Vero cells were cultured in full growth medium (cell culture media), that is, minimum Eagle's medium (MEM) supplemented with 10% fetal bovine serum (FBS, HyClone) and 1% penicillin-streptomycin (PS) solution (x100 HyClone). U2OS cells were cultured in full growth medium (cell culture media), that is, McCoy's 5A(PM150710) supplemented with 10% fetal bovine serum (FBS, HyClone) and 1% penicillin-streptomycin (PS) solution (x100 HyClone). The culturing condition was humidified atmosphere at 37 ℃ charged with 5% $CO_2$.

HeLa cells were transient transfected with Halo-Sec61β (Addgene plasmid # 123285), using Lipofectamine™ 3000 reagent following standard protocol. U2OS cells were transient transfected with Halo-H2B, Halo-TOMM20 (Addgene plasmid # 123284) using Lipofectamine™ 3000 reagent following standard protocol. The transfected cells were diluted by the stable sieve with G418 drug to obtain a single cell line with stable Halo- Sec61β, Halo-H2B, Halo-TOMM20 expression.

Vero cells were transient transfected with Halo-βTubulin (Addgene plasmid # 64691) using Lipofectamine™ 3000 reagent following standard protocol.

4.2 Total internal reflection microscopy

Super-resolution imaging was studied with a total internal reflection fluorescence microscope (TIRFM) built on an Olympus IX71 inverted microscope as described earlier. The laser light was focused on the back focal plane of a × 100 objective (UAPON 100XOTIRF; 1.49 numerical aperture). An EMCCD camera (iXon DU-897U) was implemented for data acquisition.

4.3 Super-resolution imaging acquisition

Halo-H2B expressing cells were incubated with 500 nM STMR/SRhB for 1 h. Halo-Sec61β and Halo-βTubulin expressing cells were incubated with 300 nM STMR/SRhB for 2 h. Halo-TOMM20 expressing cells were incubated with 50 nM STMR for 2 h. The free remaining dyes were washed with PBS for three times, and were further cultured in $CO_2$ incubator with fresh MEM media for 30 min. The imaging media was MEM without phenol red supplemented with 10% FBS.

A conventional image was acquired with low laser intensity before super-resolution imaging. During super-resolution imaging, a continual 532 nm laser (2 kW/cm$^2$) was utilized for excitation. The single-molecule photoswitching signals were recorded at 200 Hz for Halo-Sec61β, Halo-TOMM20 expressing cells and 25 Hz for Halo-H2B, Halo-βTubulin expressing cells.

4.4 Post-Processing of Super-resolution Imaging Data

Super-resolution imaging analysis was performed in either a ThunderStorm plugin[5] of ImageJ[6] and a home-written Matlab software. Briefly, the raw frames were filtered with a difference-of-Gaussians filter to search for signal candidates. Then the point spread functions (PSF) of those candidates were fitted with an integrated form of symmetric 2D Gaussian function (Fitting radius: 3.0 pixel) following Maximum likelihood method to estimate the precise location and single-molecule intensity. The localization precision was calculated according to the Thompson formula.[7] Those PSFs with large widths (> 1.5×median(sigma)), small widths (< 0.5×median(sigma)) were eliminated.

Camera readout intensity (I) was converted to photons through the below equations:

$$\text{photons} = \frac{\text{I}\times\text{ADU}}{\text{QE}\times\text{EMGain}} \tag{2}$$

I was the intensity value direct read from camera. ADU was the sensitivity of EMCCD (15.82 electrons per A/D count). EMGain was the gain configuration of the camera (100 in this study). QE was the photon efficiency of the camera.

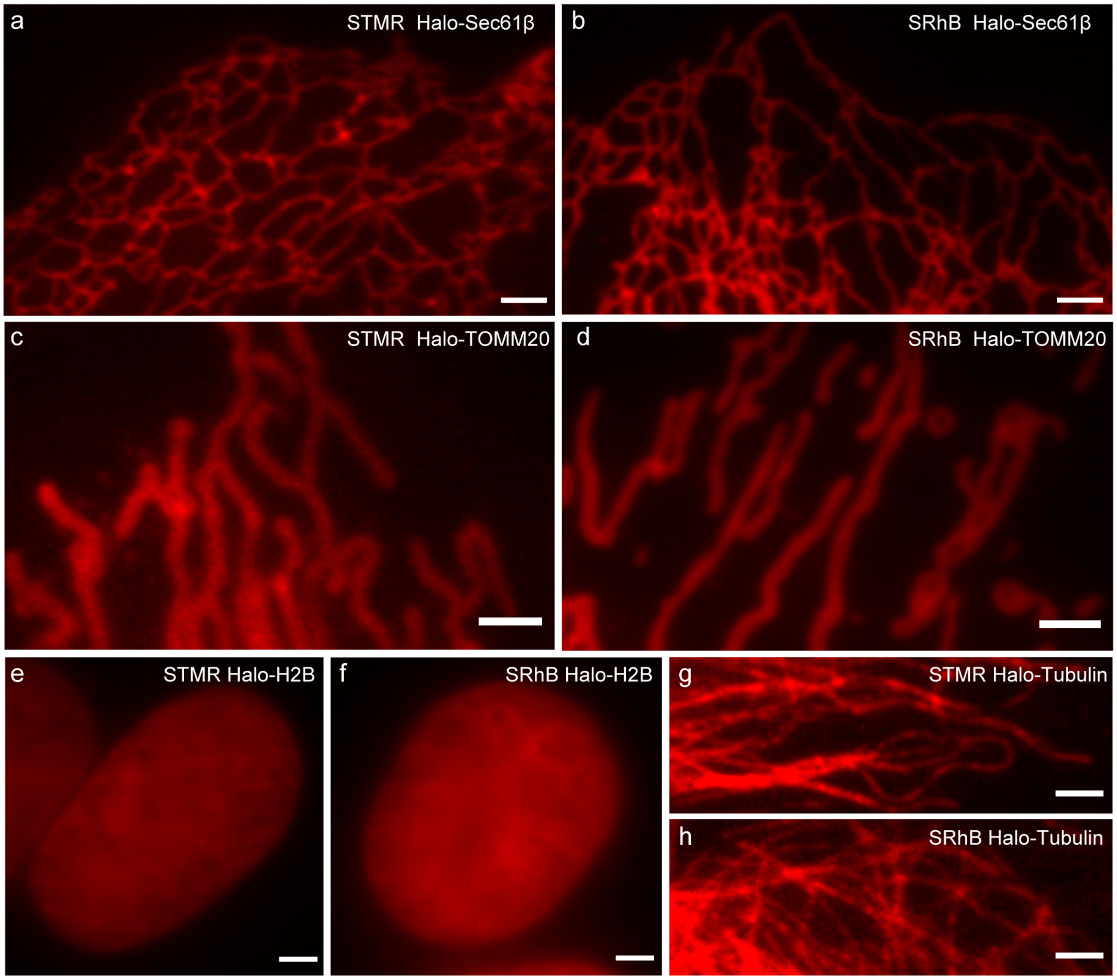

**Figure S3**. The conventional image of different target fusion proteins in living cells stained with STMR and SRhB. Scale bars: 2 μm (a-e); 1 μm (g, h).

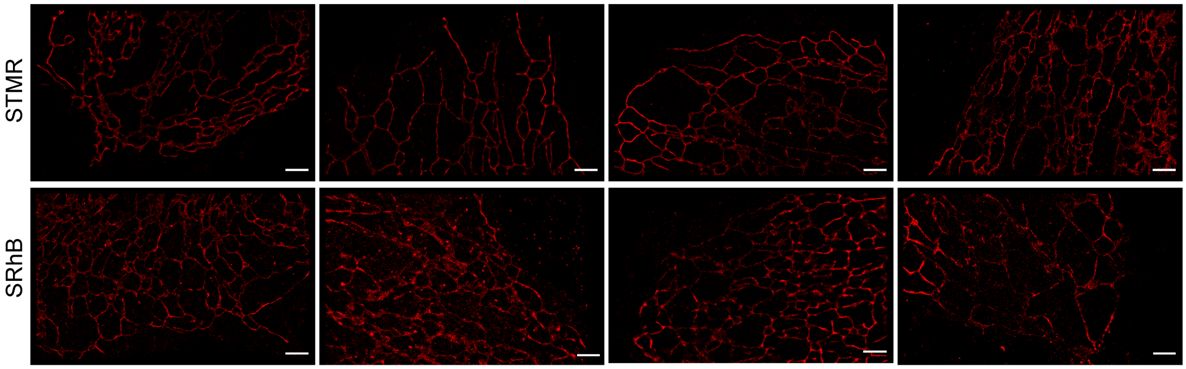

**Figure S4**. Comparison on the super-resolution imaging of endoplasmic reticulum in living cells stained with STMR and SRhB. Scale bars: 2 μm.

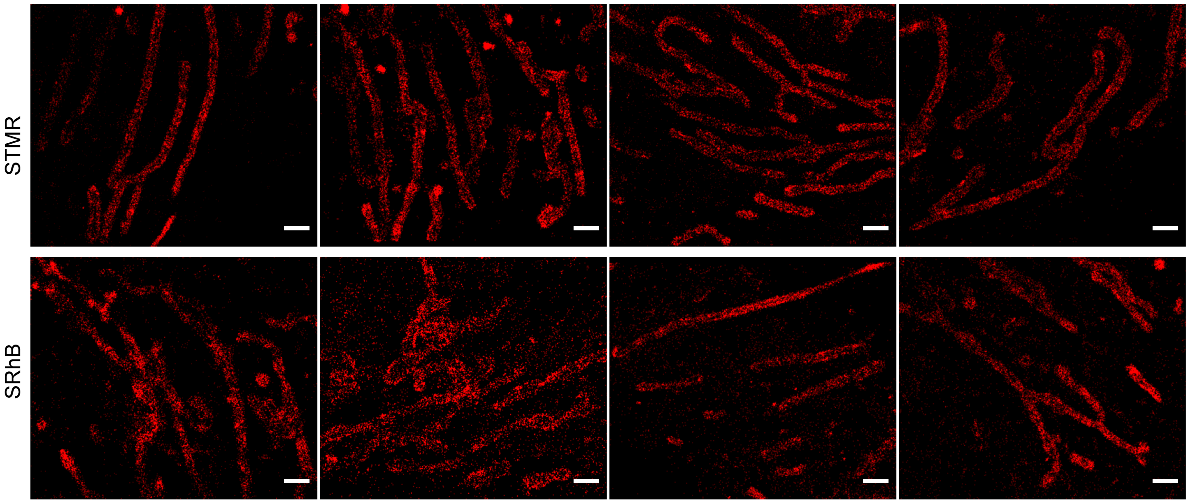

**Figure S5**. Comparison on the super-resolution imaging of mitochondria in living cells stained with STMR and SRhB. Scale bars: 2 μm.

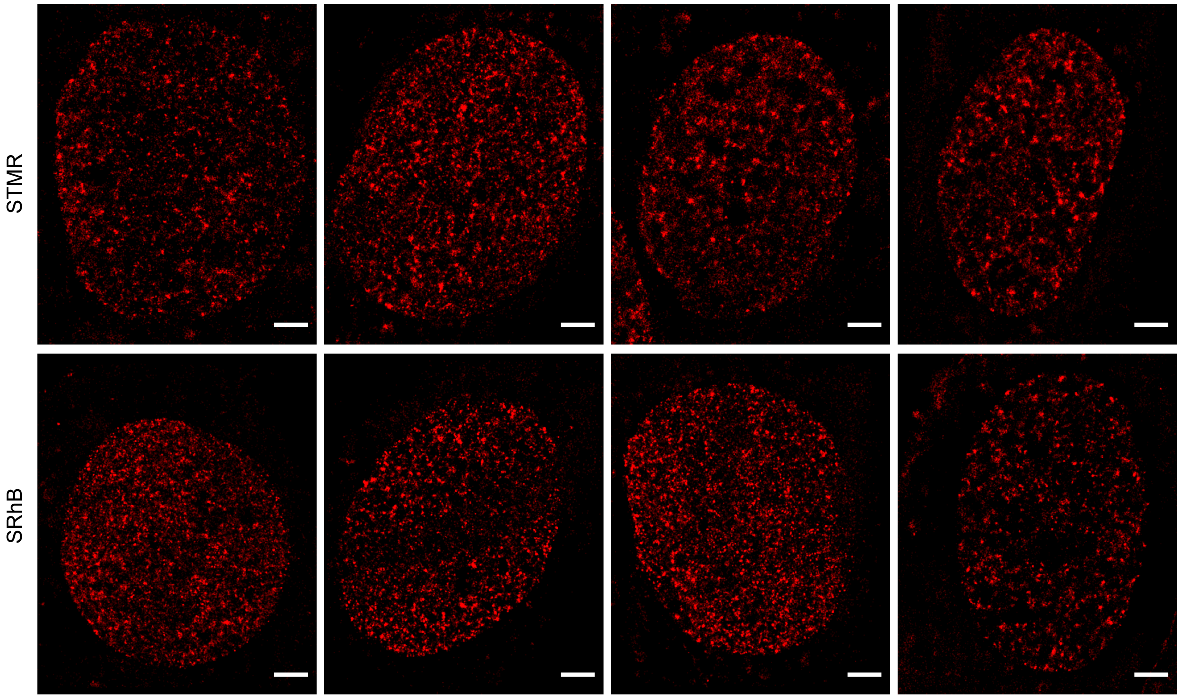

**Figure S6**. Comparison on the super-resolution imaging of H2B protein in living cells stained with STMR and SRhB. Scale bars: 2 μm.

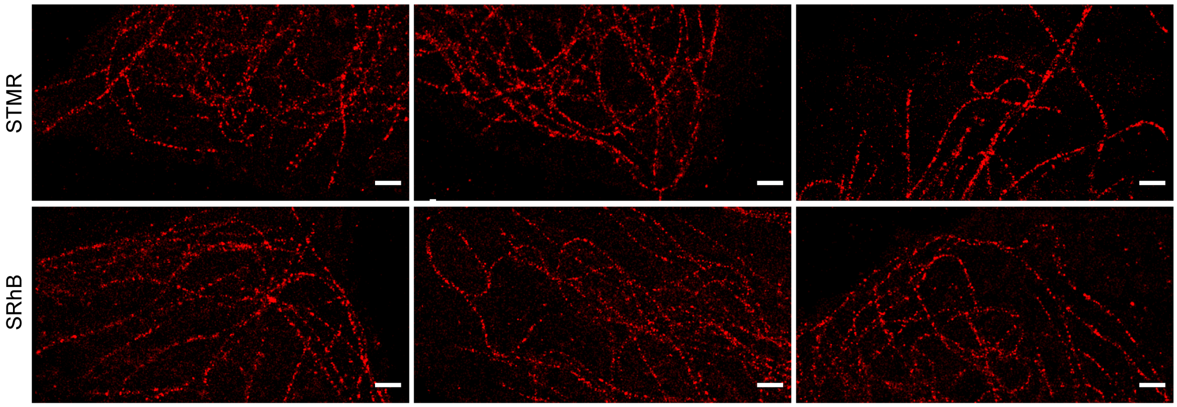

**Figure S7**. Comparison on the super-resolution imaging of microtubules in living cells stained with STMR and SRhB. Scale bars: 2 μm.

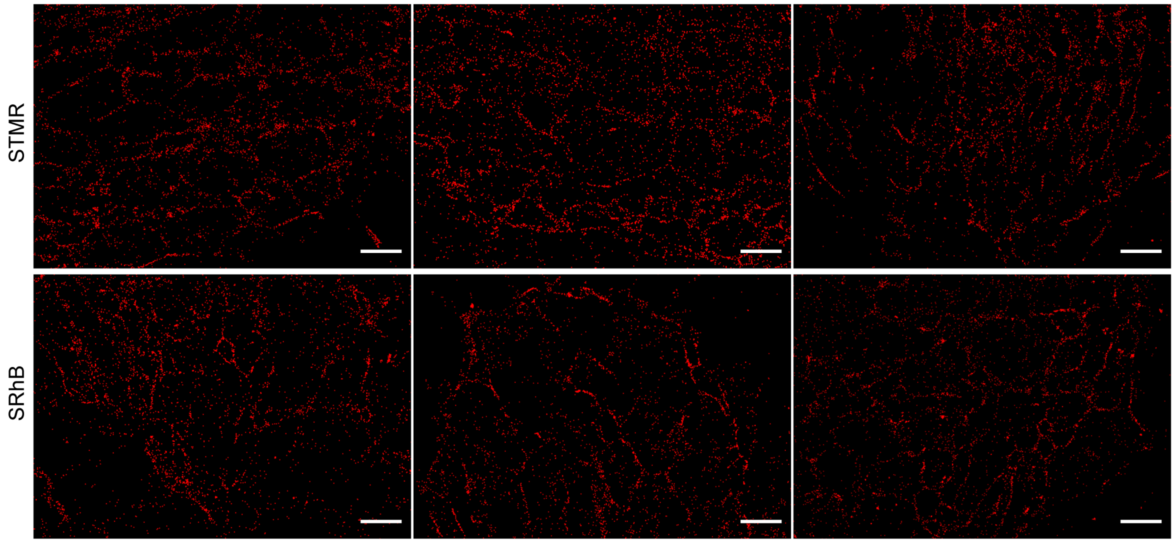

**Figure S8**. The super-resolution imaging of endoplasmic reticulum in living cells stained with STMR and SRhB under 40 ms exposure time per frame. Scale bars: 2 μm.

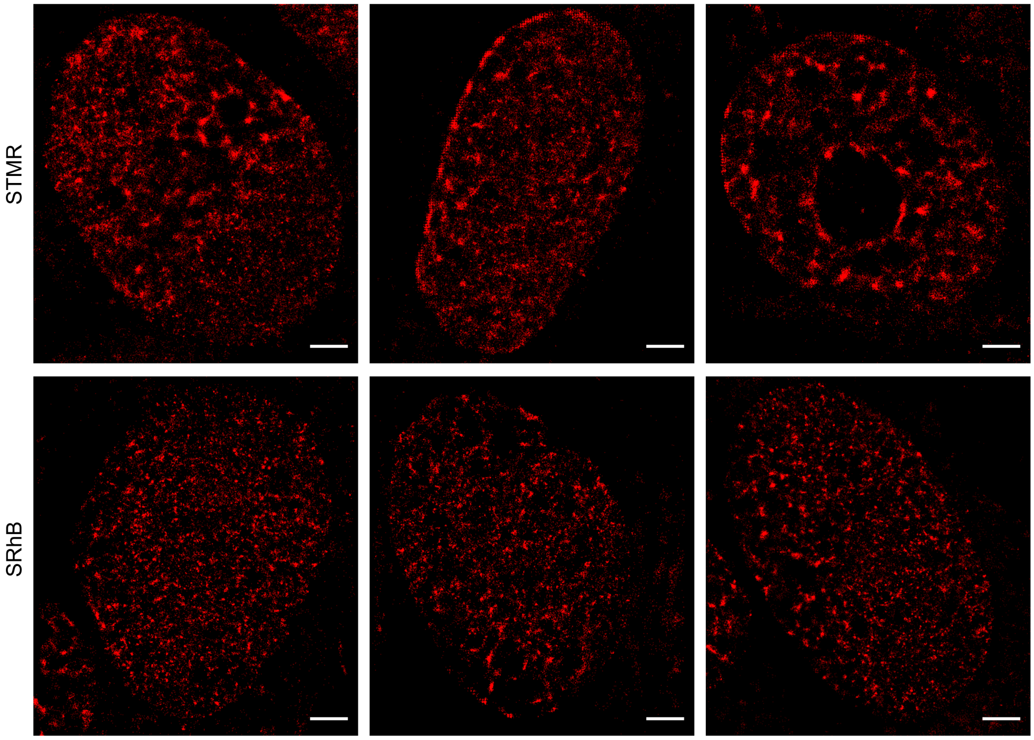

**Figure S9**. The super-resolution imaging of H2B protein in living cells stained with STMR and SRhB under 10 ms exposure time per frame. Scale bars: 2 μm.

### 5 Characterization spectra

5.1 [1]H NMR spectrum of SRhB-COOH

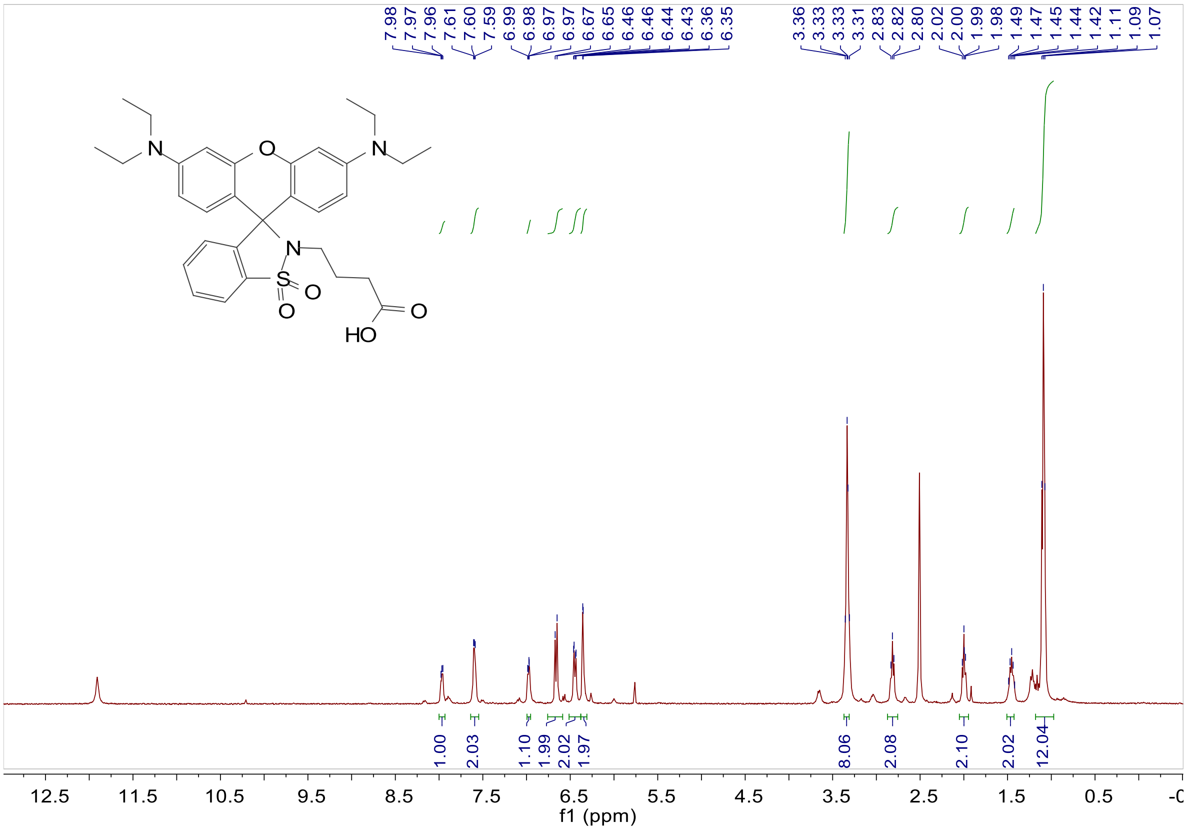

5.2 [13]C NMR spectrum of SRhB-COOH

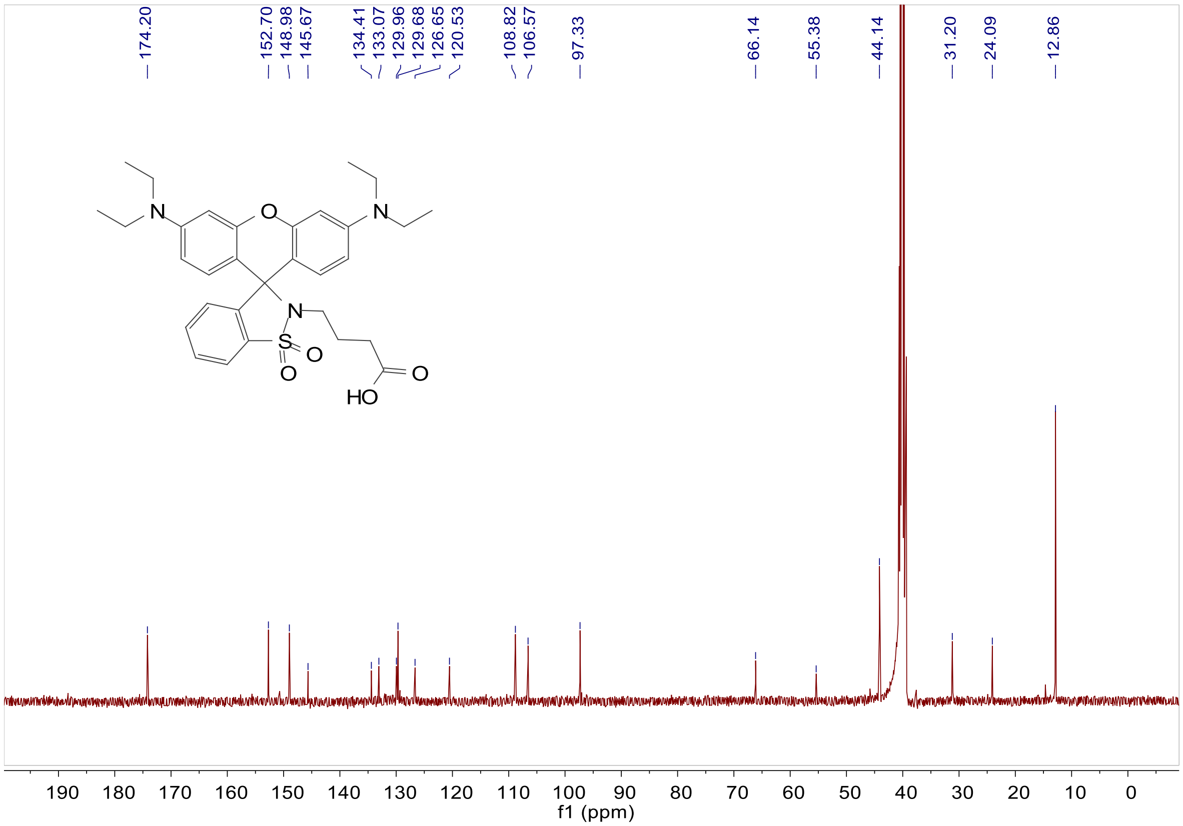

5.3 $^{1}$H NMR spectrum of SRhB

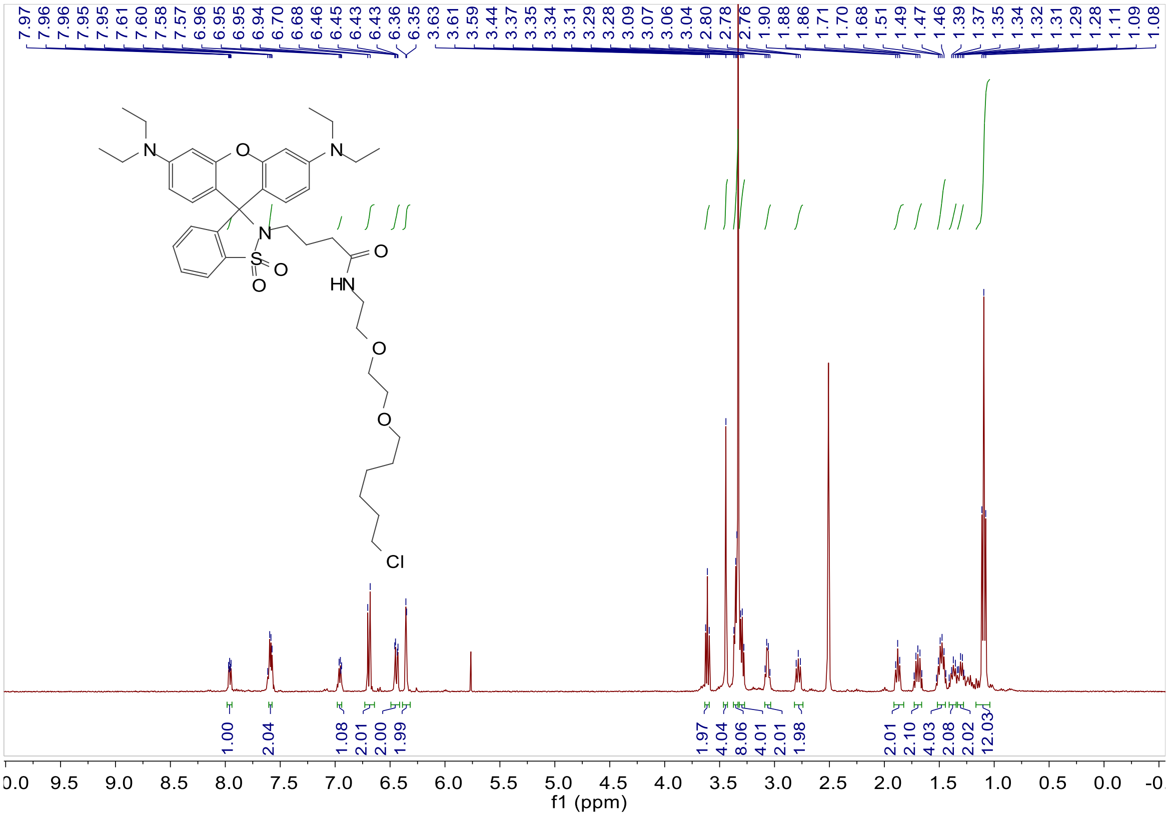

5.4 $^{13}$C NMR spectrum of SRhB

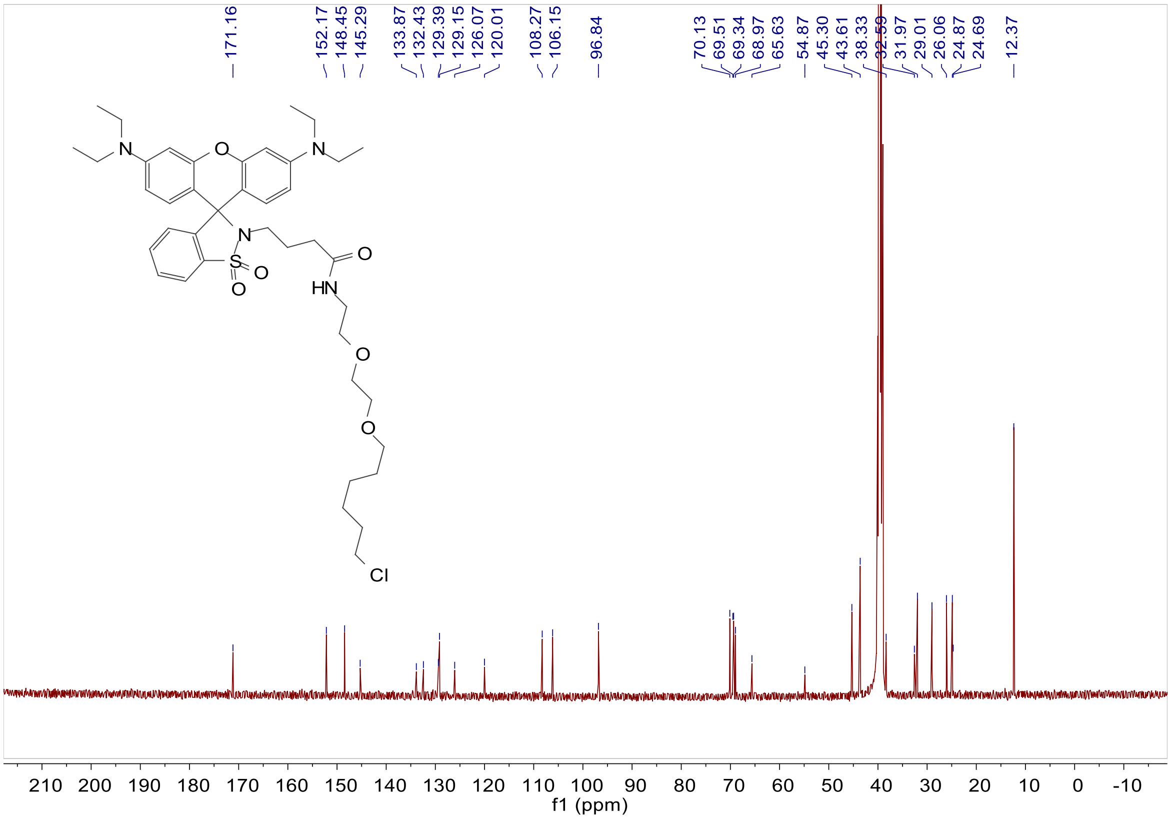

### 6 Reference

1. Ye, Z.; Yu, H.; Yang, W.; Zheng, Y.; Li, N.; Bian, H.; Wang, Z.; Liu, Q.; Song, Y.; Zhang, M.; Xiao, Y., Strategy to Lengthen the On-Time of Photochromic Rhodamine Spirolactam for Super-resolution Photoactivated Localization Microscopy. *J. Am. Chem. Soc.* **2019,** *141* (16), 6527-6536.
2. Ye, Z.; Yang, W.; Wang, C.; Zheng, Y.; Chi, W.; Liu, X.; Huang, Z.; Li, X.; Xiao, Y., Quaternary Piperazine-Substituted Rhodamines with Enhanced Brightness for Super-Resolution Imaging. *J. Am. Chem. Soc.* **2019,** *141* (37), 14491-14495.
3. Zheng, Y.; Ye, Z.; Liu, Z.; Yang, W.; Zhang, X.; Yang, Y.; Xiao, Y., Nitroso-Caged Rhodamine: A Superior Green Light-Activatable Fluorophore for Single-Molecule Localization Super-Resolution Imaging. *Anal. Chem.* **2021,** *93* (22), 7833-7842.
4. Ye, Z.; Zheng, Y.; Peng, X.; Xiao, Y., Surpassing the Background Barrier for Multidimensional Single-Molecule Localization Super-Resolution Imaging: A Case of Lysosome-Exclusively Turn-on Probe. *Anal. Chem.* **2022,** *94* (22), 7990-7995.
5. Ovesný, M.; Křížek, P.; Borkovec, J.; Švindrych, Z.; Hagen, G. M., ThunderSTORM: a comprehensive ImageJ plug-in for PALM and STORM data analysis and super-resolution imaging. *Bioinformatics* **2014,** *30* (16), 2389-2390.
6. Schneider, C. A.; Rasband, W. S.; Eliceiri, K. W., NIH Image to ImageJ: 25 years of image analysis. *Nat. Methods* **2012,** *9* (7), 671-675.
7. Thompson, R. E.; Larson, D. R.; Webb, W. W., Precise nanometer localization analysis for individual fluorescent probes. *Biophysical journal* **2002,** *82* (5), 2775-83.
